# Adaptive HD-sEMG decomposition: Towards robust real-time decoding of neural drive

**DOI:** 10.1101/2023.09.18.558259

**Authors:** Dennis Yeung, Francesco Negro, Ivan Vujaklija

**Affiliations:** Department of Electrical Engineering and Automation, Aalto University,Espoo, Finland; Department of Clinical and Experimental Sciences, Università degli Studi di Brescia, Brescia, Italy

## Abstract

Neural interfacing via decomposition of high-density surface electromyography (HD-sEMG) should be robust to signal non-stationarities incurred by changes in joint pose and contraction intensity. We present an adaptive real-time motor unit (MU) decoding algorithm and test it on HD-sEMG collected from the extensor carpi radialis brevis during isometric contractions over a range of wrist angles and contraction intensities. The performance of the algorithm was verified using high-confidence benchmark decompositions derived from concurrently recorded intramuscular electromyography (iEMG). In trials where contraction conditions between the initialization and testing data differed, the adaptive decoding algorithm maintained significantly higher decoding accuracies when compared to static decoding methods. Using ‘gold standard’ verification techniques, we demonstrate the limitations of filter re-use decoding methods and show the necessity of parameter adaptation to achieve robust neural decoding.

## 1. Introduction

Force generation in human skeletal muscles is governed by the activity of constituent motor units (MUs). Each MU is comprised of a single alpha motor neuron and the set of muscle fibers that it innervates, where a single axonal action potential initiates a tension-generating contractile twitch in the innervated fibers. The discharge pattern of a MU population thus encode the neural drive underlying gross muscular contraction [1,2]. Historically, the precise activation times of individual MUs were only attainable via manual or semi-automatic spike sorting of electromyography (EMG) signals measured from indwelling electrodes [1, 3–6]. More recently, convolutive blind source separation techniques have been developed to automatically extract MU spike trains from high-density surface EMG (HD-sEMG) [7–9]. Such methods yield detailed neural information in a non-invasive manner and are capable of extracting far more MUs compared to the spike sorting of intramuscular EMG (iEMG) [10]. For these reasons, HD-sEMG decomposition has garnered considerable interest in studies on neurophysiology, motor control and neuromuscular disorders [10–13]. In particular, MU decomposition offers practical advantages over established modes of human-machine interfacing (HMI) due to the access to higher neural information without the need of invasive procedures [14–16]. Traditionally, decomposition yields a set of separation vectors (MU filters) that distill HD-sEMG into underlying source activities. This process relies on repeated execution of iterative numerical methods over observations spanning substantial periods of time, typically 10 s or more [8, 17]. Hence, such batch decomposition algorithms are unsuitable for realtime deployment. Instead, reapplication of batch-decomposed MU filters to real-time measurements has been a commonly adopted approach [16, 18, 19]. However, these techniques assume surface MU action potentials (sMUAPs) to remain consistent. In reality, factors such as fatigue, contraction intensity, and joint position alter the expression of sMUAPs on the skin surface [20–23]. To tackle this challenge, decoding algorithms that adapt to new data have been developed [17, 24]. However, these methods have been tailored to specific conditions and are yet to be evaluated against the gold standard reference of iEMG-decomposed spike trains.

Here we propose a real-time MU decoding algorithm that updates the MU filter and signal preprocessing transforms as new action potentials of the observed MU emerge. The algorithm was evaluated on HD-sEMG recordings pertaining to isometric wrist extension contractions that vary across contraction intensities and joint angles. The accuracy of the algorithm was verified using reference spike trains manually decomposed from concurrently recorded finewire iEMG.

## 2. Methods

### 2.1 HD-sEMG decomposition

The decomposition techniques employed in this work are based on a convolutive mixture model for EMG generation:

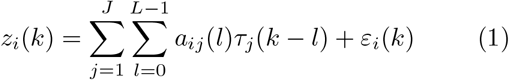

where *z*_*i*_(*k*) is the value of HD-sEMG channel *i* at time instant *k. τ*_*j*_(*k − l*) is the pulse train of MU *j* while *a*_*ij*_(*l*) encodes its respective action potential. *L* is therefore the maximum duration of impulse responses that is considered in the model and *ε*_*i*_(*k*) is additive noise inclusive of the activities of unextractable MUs.

The algorithm for batch decomposition is described in detail in [8] though, here, a brief overview will be given for completeness. Firstly, the HD-sEMG signals are extended by appending their time-delayed versions as additional observations. This conditions the data for the FastICA algorithm [25], which normally decomposes instantaneous mixtures, to handle convolutive mixtures [26]. Further preprocessing of the observations includes zero-phase component sphering, which aids in the convergence of FastICA [26]. The batch algorithm then extracts underlying source activities in a sequential manner, thereby estimating the firing intervals of MUs responsible for the generation of the observed HD-sEMG. Each source signal is extracted as:

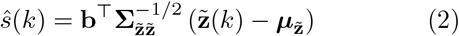

where 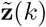 is the extended observation vector and 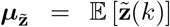 is the vector of subtractive means for centering 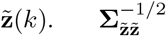 is the sphering matrix, calculated as the inverse square root of the covariance matrix of extended obsrvations, 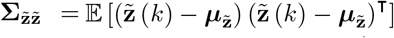 Finally, **b** is the spatiotemporal filter that extracts the MU contribution.

The processes involved for simultaneously estimating **b** and ŝ (*k*) in a blind manner can be broken down into the *Extraction* and *Refinement* step. The *Extraction* step employs FastICA which iterates a fixedpoint algorithm with an objective function optimizing the sparsity of ŝ (*k*). Orthogonalization of MU filters is performed at every iteration to ensure convergence to new sources. In the *Refinement* step, the MU filters and spike trains are iteratively updated to optimize the silhouette measure (SIL), a value which measures the accuracy of the separation [8, 23]. As per [7], each iteration first involves peak detection on the estimated source signal. Spike classification is then performed where the kmeans++ algorithm is used to distinguish peaks as either spikes or noise, with cluster centroids *c*_*hi*_ and *c*_*lo*_, respectively. Finally the MU filter is re-calculated as the cross-correlation between the sphered, extended observations and the current estimated spike train:

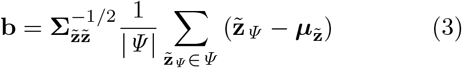

Where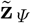 represents members in a set of extended observations corresponding to time instants of estimated spikes, 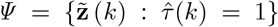The *Refinement* step is thus repeated until the SIL value of the re-estimated source ceases to increase and sources with a final SIL value above a minimum acceptance threshold are deemed as viable MU pulse trains.

### 2.2 Online decomposition

#### 2.2.1 Static decoding

So far, the most prominent approach to estimating MU activities in real-time is through the re-use of the MU filter, **b**, and preprocess transforms, and 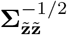 as presented by Barsakcioglu et. al. [27]. These are initially obtained from training data and then continuously reapplied to new windows of extended data in the same manner as equation (2). Detected peaks in the estimated source signal are further sorted as either spikes or noise peaks. Rather than using the kmeans++ algorithm, this is simply determined by a threshold set at the midpoint between the spike and noise centroids, *c*_*hi*_ and *c*_*lo*_, also retained from batch decomposition of the training data. To accommodate for deviations between the conditions of the training data and the new, unseen data, this decision boundary may be altered by a relaxation factor, 0 *≤ α ≤* 1:

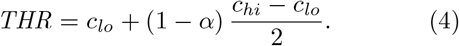

#### 2.2.2 Adaptive MU decoding

While relaxation of the decision boundary may decrease the likelihood of missed spikes, it can also lead to an increase in false positives. To address this, we propose adaptation of the MU filter and preprocess transforms as potential spike events are detected. Here, adaptation of decoding parameters occurs in parallel to the online decoding process. The updating of the MU filter is performed in a manner similar to the *Refinement* step of the batch algorithm. Since this does not rely on FastICA, re-computation of ZCA sphering transforms is unnecessary. Rather, only the inverse of the sample covariance matrix is needed. The equivalent of source extraction (equation (2)) for the adaptive decoding algorithm can therefore be written as:

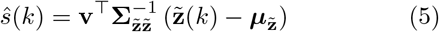

where **v** is now the MU filter and 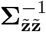 is the inverse of the observation covariance matrix.

With each new data window, temporary transforms are first derived from the updated statistics of extended observations:

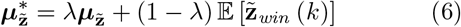

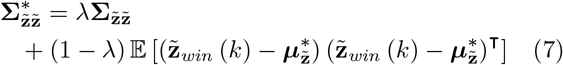

where 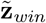 (*k*) is the *k*^th^ sample in the new window of extended data and 0*≤ λ≤* 1 is the forgetting factor that controls the influence of new data. An initial estimation of the source signal is then calculated:

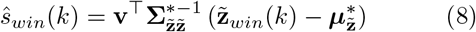

where 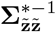 is the inverse of the temporary covariance matrix obtained from equation (7). To identify potential new spike instances for learning, the sparseness property of MU firing patterns can be leveraged. First, peak detection is conducted on the estimated source signal, ŝ_*win*_(*k*). With *a-priori* knowledge regarding the short time-span that the data window corresponds to, any strong responses to the MU filter, meaning potential spikes, will appear as outliers in the distribution of rectified peak amplitudes. Hence, rectified peak amplitudes with z-scores above a certain threshold will have their corresponding extended observation vectors added to set *Ψ*_*mem*_. Functionally, *Ψ*_*mem*_ is implemented as a first-in-first-out (FIFO) storage buffer of constant size, initialized from extended observations corresponding to spike events in the training data. As candidate spikes are detected from new data windows and new observation vectors are added to *Ψ*_*mem*_, past observations are discarded. With each update of *Ψ*_*mem*_, the MU filter is recalculated using equation (9):

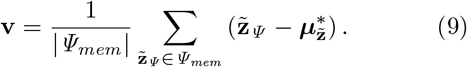

The spike and noise centroids, which are used for online spike detection outside of the adaptation algorithm, are subsequently updated. The spike centroid is recalculated via equation (10) which corresponds to the squared average amplitude of peaks extracted from the observations stored in *Ψ*_*mem*_. The noise centroid is then updated as a *λ*-weighted merging of the past *c*_*lo*_ and the average of noise peak amplitudes detected from the new source signal window:

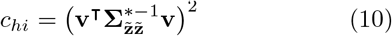

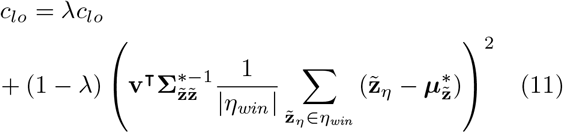

where *η*_*win*_ is the set of observations corresponding to noise peaks detected in ŝ_*win*_ (i.e. peak observations not added to *Ψ*_*mem*_).

Algorithm 1 summarizes this entire process for real-time updating of decomposition parameters. Prior to implementation, there are three static parameters that need to be defined: the threshold z-score value for accepting new observations into *Ψ*_*mem*_, the rate of forgetting, *λ*, and the cardinality of *Ψ*_*mem*_. For the results obtained in this work, the corresponding values used were 3.3, 0.985 and 110, respectively, based on initial testing.

### 2.3 Experimental setup

Five able-bodied subjects were recruited for the experiment, four male, one female, ages 29 - 34, all right-handed. The study was approved by the local ethical board of Aalto University (approval number D/505/03.04/2022). Prior to the experiments, all subjects gave their written informed consent in accordance with the Declaration of Helsinki.

Subjects were seated for the duration of the experiment with their dominant upper-limb placed in a specialized tabletop rig designed to constrain the wrist joint at various angles of extension (Figure 1). Forces generated by isometric contractions pertaining to wrist extension were measured with a load cell (TAS606, HT Sensor Technology, China) at a sampling rate of 100 Hz. Prior to the insertion of finewire electrodes, the subject’s maximum voluntary contraction (MVC) forces were measured at wrist joint angles corresponding to 0%, 12.5% and 25% of their maximal extension, with 0% relating to a neutral wrist position. MVC was calculated as the averaged maximal force from three MVC contractions of 1.5 s long with short breaks in between each contraction to prevent fatigue.

**Figure 1.**
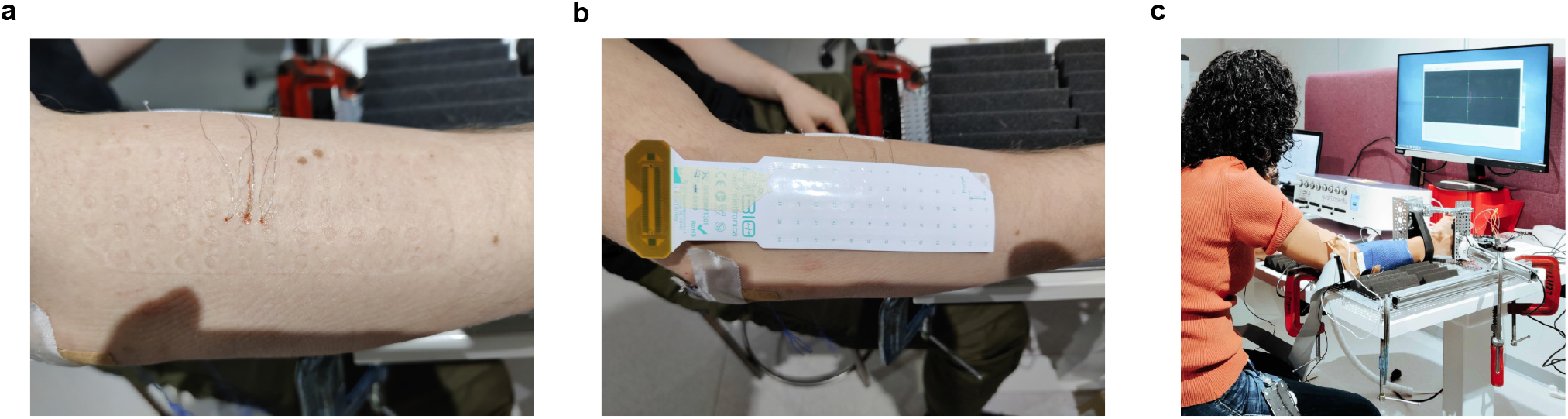
Experimental setup. (a) Three fine-wire electrode pairs inserted into a subject’s ECRB. (b) A 64-channel high-density surface electrode grid placed above the iEMG insertion sites show in (a). (c) Subject with iEMG and HD-sEMG electrodes attached to their dominant arm which has been placed inside the force measurement rig. Task cues are shown on the computer screen in front of the subject

Stainless steel/silver (SS/Ag) wires with polytetrafluoroethylene (PTFE) insulation (Spes Medica s.r.l, Italy) were used as intramuscular electrodes. The wires had a diameter of 0.11 mm with the final 3-5 mm of the recording tips stripped of the insulating material. Three insertion points were targeted, centered at the bulk of the extensor carpi radialis brevis (ECRB) and aligned down the muscle axis at approximately 4 mm intervals. Location of the ECRB was guided by [28] and palpation during wrist extension and radial deviation movements. The fine-wires were inserted as pairs (bipolar configuration) using 25G cannulae to a depth targeting MUs proximal to the skin surface. Signal inspection was conducted after the insertion of each electrode pair. If the signal was invalid (short-circuited, excessive noise, low selectivity or no viable units detected) and could not be remedied by light manipulation of the fine-wires, the wires were removed and another insertion of new electrodes was made slightly lateral to the original insertion point. The maximum

#### Algorithm 1 Adaptation of online MU decoding parameters

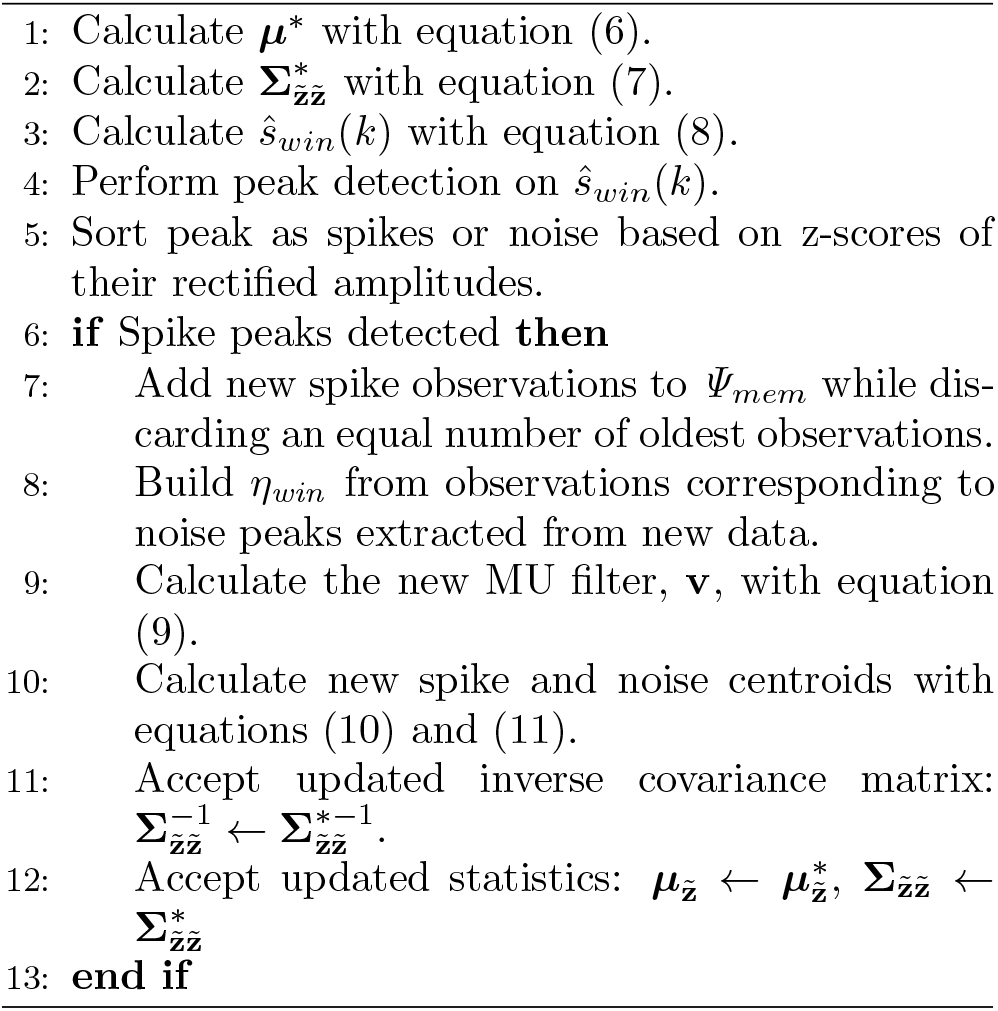

overall number of insertion attempts was bounded to five for the sake of subject comfort. The experiment only proceeded so long as at least one valid iEMG channel was attained. The bipolar iEMG signals were preamplified by an adapter (ADx5JN, OT Bioelettronica, Italy) with a gain of 5, and acquired by a bioamplifier (Quattrocento, OT Bioelettronica, Italy) with a fixed gain of 150 at 10240 Hz with 10-4400 Hz hardware bandpass filtering. Subsequent processing of iEMG signals included high-pass filtering with a 250 Hz cut-off to lower baseline noise and to produce sharper action potentials [5, 29].

Placement of the overlaying HD-sEMG matrix was conducted approximately 8 minutes after the final finewire insertion. This allowed for sufficient coagulation and minimizing the leakage of blood or plasma to the surface recording site. A 64-channel rectangular electrode matrix (GR08MM1305, OT Bioelettronica, Italy) with 8 mm inter-electrode distance was placed on top of the ECRB, centered above the iEMG insertion sites (Figure 1). Two reference electrodes (Neuroline 720, Ambu A/S, Denmark), one for the pre-amplifier and one for the bioamplifier, were placed at the medial epicondyle and olecranon process. The HD-sEMG signals were buffered by a pre-amplifier (AD64F, OT Bioelettronica, Italy) prior to being acquired by the same benchtop amplifier used for iEMG at 150 gain, 10240 Hz with 10-4400 Hz hardware bandpass filtering. Pre-processing of the HD-sEMG signals for automatic decomposition included downsampling to 2048 Hz and bandpass filtering with 10-900 Hz cut-offs.

Prior to the commencement of recordings, subjects were asked to perform slow dynamic wrist extensions, up to 25% of maximum range of movement, to allow the settling-in of fine-wire electrodes and HD-sEMG matrix. The recording and cueing of contractions were facilitated by a custom Matlab R2021b (MathWorks Inc., USA) framework. All subject cues, along with the real-time force feedback, were displayed on a computer screen.

### 2.4 Experimental protocol

Isometric wrist extension contractions with trapezoidal force profiles (5 s ramp, 20 s plateau) were recorded at different joint angles and different force levels. For subjects A and B, contractions were recorded at force levels of 5%, 10% and 15% MVC at wrist joint angles of 0% and 25% maximal extension. For subjects C, D and E, contractions of 5%, 7.5% and 10% MVC were recorded at 0%, 12.5% and 25% maximal wrist extension. Recordings progressed from 0% to 25% extension while the order of force levels recorded was randomized. Three repetitions were recorded for each contraction condition.

### 2.5 Obtaining iEMG decomposition benchmarks

#### 2.5.1 Extraction of MU activity concurrent in iEMG and sEMG

To identify MUs present in both surface and intramuscular signals, a two-stage semi-automatic technique was employed. For each repetition, a set of MUs and their respective spike trains are first extracted from HD-sEMG via the batch decomposition method described in Section 2.1. The resultant spike intervals were then used to trigger action potentials in the iEMG signals. Here, MUs whose activities are present in both the concurrently recorded HD-sEMG and iEMG signals will trigger distinct intramuscular motor unit action potentials (iMUAPs). Typically, these are mono and polyphasic waveforms with peaks well above the baseline noise [4]. On the other hand, MUs that were only extractable via HD-sEMG decomposition will trigger flat iMUAPs (peak-to-peak amplitudes *<* 2*µV*). In this way, units present in both surface and intramuscular recordings are identified. The spiketrains of such MUs were then imported to EMGlab [29], a Matlab-based spike annotation software, for manual correction by an experienced operator such that a highconfidence benchmark is obtained.

#### 2.5.2 Tracking MUs across contraction conditions

MUs were matched by the same experienced operator through visual comparison of their multi-channel iMUAPs. As each iEMG channel consisted of a bipolar measurement, activity from a single source manifests as action potentials that vary greatly in profile across channels but are, albeit, time-locked. Thus, a single

MU may be characterized by up to three distinct action potentials triggered by the same spike instances. Examples are shown in Figure 2 where such iMUAP profiles may be used to manually match MUs across contraction conditions.

**Figure 2.**
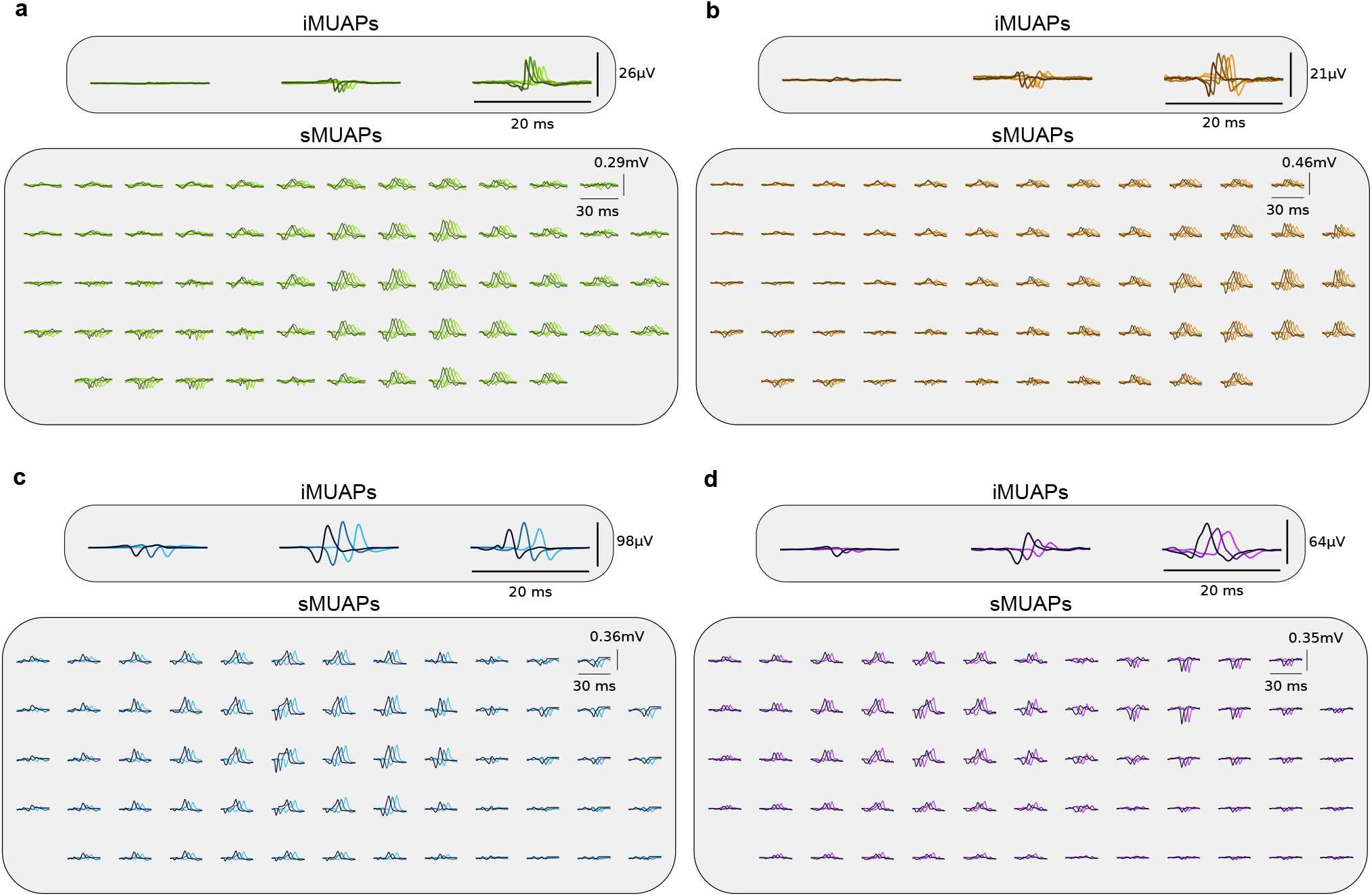
iMUAPs and sMUAPs extracted from different contraction conditions. Up to 3 unique iMUAP shapes are utilized for manual matching of MUs extracted from different contraction conditions. While each individual iMUAP profile is still susceptible to change across the angle and force conditions, causing potential matching ambiguities, the presence of multiple time-locked and distinct profiles facilitate matching of MUs that may share similar iMUAP profiles in one particular channel. Variation in the sMUAP profiles across contractoin conditions is also observed, resulting in sub-optimal extraction of source activities when using static decoding algorithms. (a) iMUAPs and sMUAPs of MU B1 extracted from all 5 contraction conditions that it was detected in. From darkest to lightest plot lines, the displayed MUAPs correspond to angle/force combinations of 0%/5%, 0%/10%, 25%/5%, 25%/10% and 25%/15%, respectively. (b) iMUAPs and sMUAPs of MU B2 obtained from the same contraction conditions as displayed in (a). (c) iMUAPs and sMUAPs of MU C1 extracted from 3 different contraction conditions. From darkest to lightest plot lines, the displayed MUAPs correspond to angle/force combinations of 0%/5%, 12.5%/7.5% and 25%/10%, respectively. (d) iMUAPs and sMUAPs of MU C2 obtained from the same contraction conditions as displayed in (c).

### 2.6 Pseudo-online testing

In the pseudo-online tests, multiple trials were conducted to gauge the robustness of the proposed adaptive MU decoding algorithm across different contraction conditions. In each trial, the decoding algorithm was initialized from one repetition and then applied to extract MU activity in another. Here, data was fed in windows of 200 ms and in time increments of 100 ms, thereby simulating real-time deployment. For comparative purposes, the static decoding technique (Section 2.2.1) was also tested using different spike threshold relaxation values, from *α* = 0 to *α* = 0.5 in increments of 0.1.

Since the decoding algorithms were to be compared in scenarios where the conditions of the training data differed from those of the test data, only MUs with high-confidence iEMG-decomposed benchmarks (obtained by methods described in Sections 2.5.1 and 2.5.2) in at least two force levels for at least two angle conditions were selected for this analysis. Table 1 lists the MUs selected for this testing along with the contraction conditions in which they were detected in. For each eligible MU, all pair-wise combinations of training and testing repetitions were analyzed.

**Table 1.** Catalog of MUs and trial conditions used for this study.

| Angle (%) | Force (%) | MU |  |  |  |  |  |  |  |  |  |  |  |  |  |  |  |  |  |
| --- | --- | --- | --- | --- | --- | --- | --- | --- | --- | --- | --- | --- | --- | --- | --- | --- | --- | --- | --- |
|  |  | A1 | A2 | B1 | B2 | B3 | C1 | C2 | C3 | C4 | C5 | D1 | D2 | E1 | E2 | E3 | E4 | E5 | E6 |
| 0 | 5 | o | o | o | o | o | o | o | o | o | o | o | x | o | o | o | o | o | x |
|  | 7.5 | - | - | - | - | - | o | o | o | o | o | o | x | o | o | o | o | o | o |
|  | 10 | o | o | o | o | o | o | o | o | x | o | o | x | o | o | o | x | x | o |
|  | 15 | o | x | x | x | x | - | - | - | - | - | - | - | - | - | - | - | - | - |
| 12.5 | 5 | - | - | - | - | - | o | o | o | o | o | x | x | o | o | o | o | o | o |
|  | 7.5 | - | - | - | - | - | o | o | o | o | o | o | o | o | o | o | o | o | o |
|  | 10 | - | - | - | - | - | o | o | o | o | o | o | o | o | o | o | o | o | o |
|  | 15 | - | - | - | - | - | - | - | - | - | - | - | - | - | - | - | - | - | - |
| 25 | 5 | o | o | o | o | o | o | o | o | o | x | x | x | o | o | x | o | o | x |
|  | 7.5 | - | - | - | - | - | o | o | o | o | x | o | o | o | o | o | o | o | o |
|  | 10 | o | o | o | o | o | o | o | x | o | x | o | o | o | o | o | o | o | o |
|  | 15 | o | o | o | o | o | - | - | - | - | - | - | - | - | - | - | - | - | - |

o = MU concurrently detected in HD-sEMG and iEMG, x = MU not concurrently detected, - = trial not recorded for subject.

For each trial, the estimated spike train was compared to the iEMG-decomposed spike train using the Rate-of-Agreement (RoA) metric:

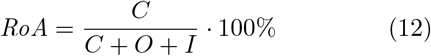

where *O* is the number of spikes that were only detected by the online decomposition algorithm and *I* is the number of firing instances exclusive to the iEMG decomposition while *C* is the number of spikes that were identified in both estimations of MU activity. In addition, two metrics that are analogous to False Negative Rate (FNR) and False Discovery Rate (FDR), when considering the iEMG-decomposed spike train as ground truth, were calculated for each trial:

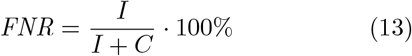

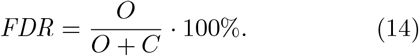

### 2.7 Statistical analysis

To detect statistically significant differences between the decoder performances, repeated-measures analysis of variance (RM-ANOVA) was conducted on the RoA values obtained from pseudo-online testing. The normality of the results was verified with Shapiro-Wilks’s testing while the assumption of sphericity was tested with Mauchly’s test. In cases where the sphericity assumption was not satisfied, Greenhouse-Geisser correction was applied to the RM-ANOVA. If the choice of decoder was found to have a significant effect, post-hoc pair-wise comparisons (Tukey-Kramer) were conducted. In all analyses, significance levels of 0.05 were used.

In addition to analyzing the full set of results, three auxiliary analyses, “Intra-condition”, “Inter-angle” and “Inter-force”, were conducted on different subsets of the data. In the Intra-condition analysis, only the trials where the training and testing data had identical angle/force conditions were considered.

In the Inter-angle analysis, trials where the training and testing data were recorded from identical force conditions, but had different angle conditions, were considered. Finally, the Inter-force analysis focused on trials where the training and test recordings consisted of contractions with identical angle but different force conditions.

## 3. Results

Figure 3 shows the results from pseudo-online testing of the static and adaptive decoders in terms of RoA with iEMG-referenced benchmark decompositions. Statistically significant differences between decoder performances were detected in all analyses (*F* (1.99, 1846.33) = 325.95, *p <* 0.001, *F* (2.16, 265.67) = 101.88, *p <* 0.001, *F* (1.99, 356.77) = 87.99, *p <* 0.001, *F* (2.00, 438.80) = 78.94, *p <* 0.001 for Global, Intra-condition, Inter-angle, and Inter-force, respectively). In post-hoc comparisons, the proposed adaptive decoding algorithm significantly outperformed static decoding for all tested *α* values (0-0.5) in the Global, Inter-angle, and Inter-force analyses (*p <* 0.001 for all comparisons). Overall, *α* = 0.3 gave the best static decoder performance with an RoA of 77.1% *±*25.2% in the Global analysis. Still, this was exceeded by the adaptive decoder by 6.7% *±*0.2%. Similarly, in the Inter-angle and Inter-force analyses, the best static decoding performances were 75.7% *±*25.0% (*α* = 0.3) and 80.8% *±*22.4% (*α* = 0.2) RoA, respectively. The adaptive decoder also outperformed these by 8.0% *±* 0.4% and 5.1% *±* 0.4%, respectively.

**Figure 3.**
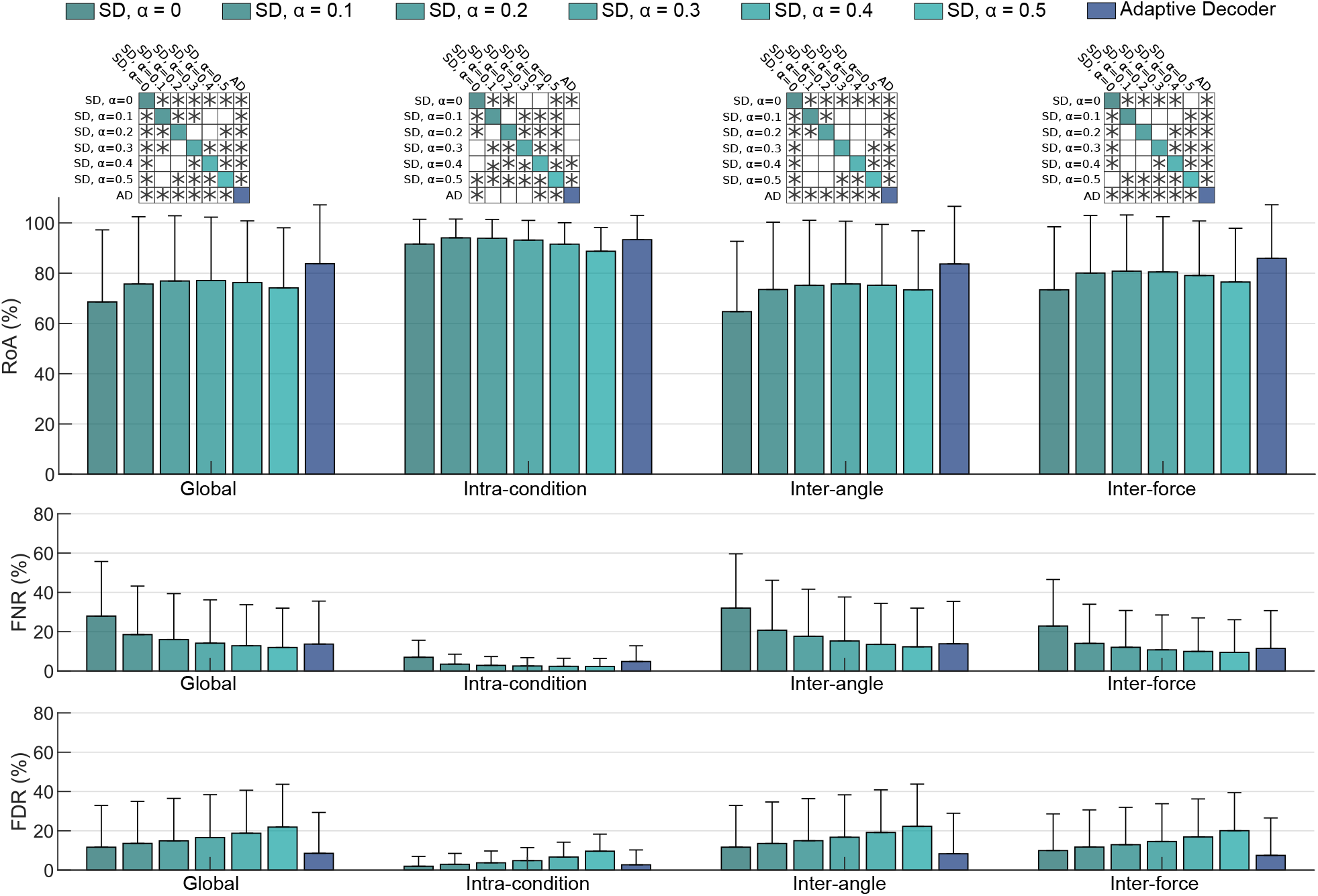
(Top) Performance of static and adaptive MU decoding (SD and AD, respectively) methods in terms of RoA with iEMGreferenced benchmark decompositions. Pairwise comparisons between decoders with statically significant differences are indicated by “*∗* “ in the grids above the bar-charts. The proposed adaptive algorithm is shown to achieve superior robustness as high RoA is maintained when tested contraction conditions differ from those used for decoder initialization. (Middle and bottom) FNR and FDR for each the decoder is also shown. Adaptive decoding is shown to negate the trade-off in increasing *α*, which lowers the occurrence of false negatives at the expense of higher false positive rates.

In the Intra-condition analysis, static decoding is shown to still perform well with *α* = 0.1 and 0.2 yielding the highest average RoAs of 94.1% *±*7.5% and 93.9%*±* 7.5%, respectively. Adaptive decoding marginally underperformed these by *−*0.7% *±*0.2% and*−* 0.6% *±*0.2%, respectively.

Figure 3 also shows decoding performances in terms of FNR and FDR. Here, the effect of increasing *α* is clearly shown. By relaxing the spike amplitude threshold, fewer spikes are missed (lower FNR) but in turn, more noise peaks are misclassified as spikes (higher FDR). In contrast, the adaptive decoding algorithm maintains low rates of either misclassification types. Compared to static decoding with *α* = 0.3, which yielded 14.2%*±* 13.7% FNR and 16.6% *±*21.8% FDR in the Global analysis, adaptive decoding achieved lower misclassifications by *−*0.5% *±*0.2% and *−*8.1% *±*0.2%, respectively. The adaptive decoding algorithm therefore resolves this trade-off between FNR and FDR. Figure 4 shows how this is achieved by comparing the source activities extracted via online decoding with static and adapting parameters. By updating the MU filtering parameters as new data is received, a clear separation of spike peaks and noise peaks in the extracted source is maintained.

**Figure 4.**
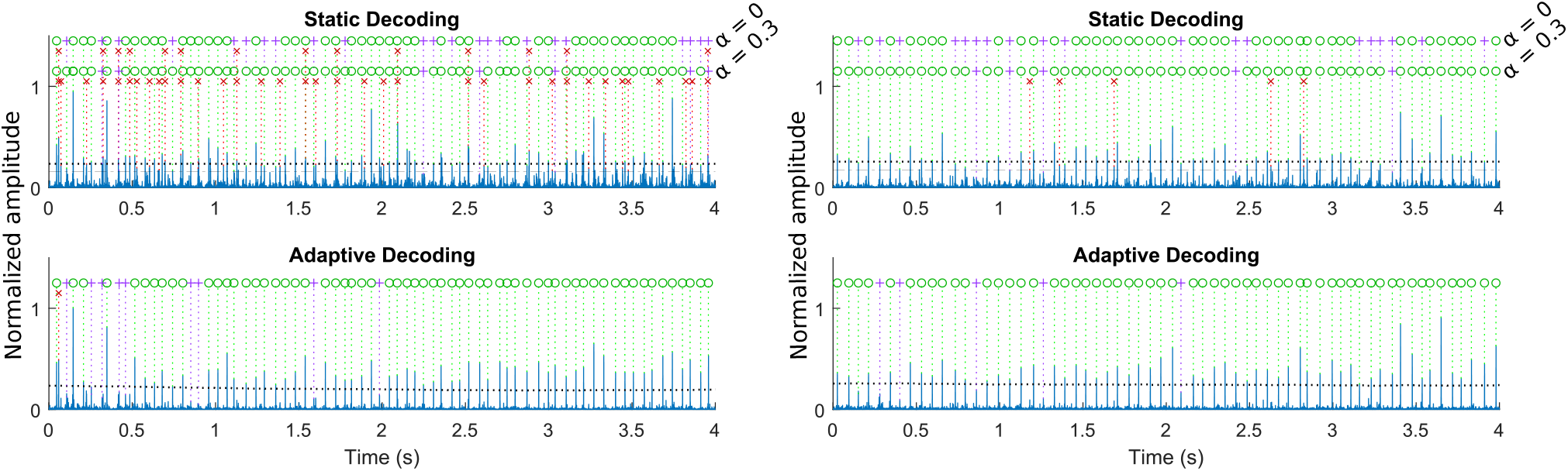
Estimated source signals and spike trains of MU E1 using static (top row) and adaptive (bottom row) decoders. Only extraction of the first 4 s out of the full 12 s recordings are shown. Estimated spikes that agree with the corresponding iEMG-referenced decomposition are indicated by green circles. Spike estimates that disagree are indicated by red x’s while missed spikes are indicated by purple crosses. For static decoders, spike estimations using *α* = 0 and 0.3 are shown. Spike amplitude thresholds are plotted as black dotted lines and dash-dot lines for the relaxed threshold (*α* = 0.3). The plots on the left side show the application of decoding parameters initialized from a wrist angle/force level combination of 0%/7% on a contraction of 25%/10%. The right side plots show results with swapped initialization and test data. With static decoding, the extracted signals are noisy and result in numerous misclassifications. The amount of missed spikes can be reduced by relaxing the spike amplitude threshold but this results in higher occurrences of misidentified spikes. With adaptive decoding, continuous updating of decomposition parameters maintain a clear separation of spike and noise peaks which result in higher decoding accuracies.

The RoA values of all trials regarding a single MU are shown in Figure 5. Results obtained via static decoding with *α* = 0 and 0.3, which yielded the best overall static decoding performance, and the proposed adaptive decoding algorithm are shown. The adaptive algorithm yielded similar decoding accuracies as static decoders in the trials that fall under the Intra-condition analysis (cells proximal to the diagonals of the heatmaps). However, in the majority of trials where the training condition does not match the test condition, adaptive decoding achieved a much higher RoA with the iEMG-decomposed benchmarks. Still, there remain cases where adaptation is unable to compensate for the large changes to the sMUAPs.

**Figure 5.**
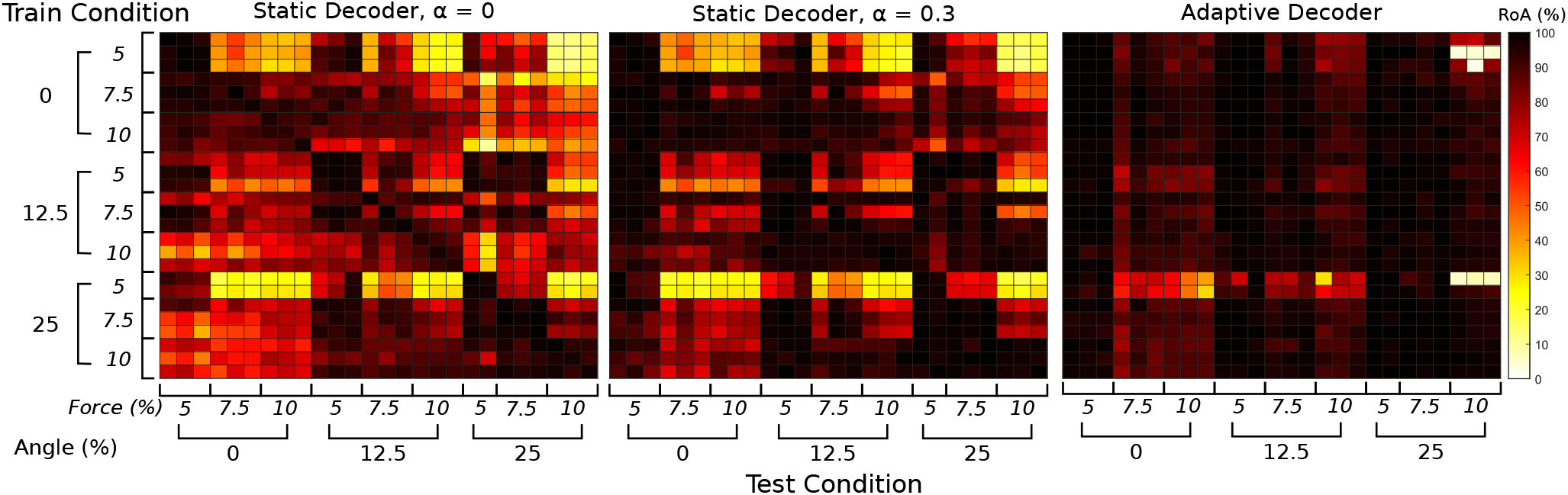
RoA results from tracking MU E1 using static and adaptive decoders. All possible pairwise combinations of training and testing repetition are shown. Cells close to the heatmap diagonals represent results used for the “Intra-condition” analysis as the decoders are tested on data pertaining to the same contraction conditions used for initialization. In such cases, static decoding with no relaxation of the spike detection threshold (*α* = 0, left) maintains high RoA with iEMG-referenced benchmarks. However, when tracking MU activity in contraction conditions that differ from the training data, RoA decreases. In some cases, this can be remedied by increasing *α* (middle), though the majority of decoding accuracies remain poor. The proposed adaptive decoder (right) therefore offers the best robustness with the majority of trials yielding RoA above 90%. However, the algorithm may not always compensate for some large changes in contraction conditions as shown by the few trials with low RoA (*<* 50%) results. For instance, the algorithm can fail to converge to an appropriate filter when presented with contractions of wrist angle/force level combinations of 25%/10% when using decoding parameters initialized from contractions of 0%/5%.

## Discussions

We have proposed an adaptive algorithm for decoding MU activity from HD-sEMG that continuously updates its internal parameters in real-time, as new measurements are acquired. Using experimental data, we demonstrate the performance of our proposed algorithm in tracking MU activities across isometric contraction conditions that vary in joint angle and intensity. In comparison to the static, non-adaptive decoding algorithm, adaptive decoding was shown to be more robust to such changes. This was verified against benchmark spike trains manually decomposed from iEMG signals that were recorded concurrently with the HD-sEMG. In terms of RoA between decoder estimations and the iEMG-referenced benchmarks, the adaptive decoder significantly outperformed static decoders across all tested spike threshold relaxation values (Figure 3). Even when the test trials only differed from the training trial by one factor (Inter-angle or Inter-force), adaptive decoding was shown to be beneficial. Nonetheless, static decoding was still effective in estimating MU activity from contractions that are similar to the training data (Intra-condition). As contraction intensity and joint angle change, so do the sMUAP profiles (Figure 2). This renders sphering transforms and MU filters derived from different contraction conditions to be sub-optimal for accurate source estimation [30, 31], resulting in missed spikes. In [23], local batch optimization of MU filters allowed for the accurate decomposition of MU activity during dynamic contractions. However, this required prior knowledge regarding the periodicity of the dynamic contractions. Here, we demonstrate that relaxation of spike acceptance thresholds can help compensate for changes to sMUAPs but this also causes an increase in false spike identifications, as evidenced by the inverse relationship between the FNR and FDR results obtained using static decoding (Figure 3). Conversely, the adaptive decoder maintains low rates of both false negative and false positive errors. This is achieved by adaptation of pre-process transforms and the MU filter as new spikes are estimated, which helps maintain a distinct separation between spike and noise peaks (Figure 4).

Previous studies on adaptive, real-time decoding algorithms include [17] and [24], both of which are based on the convolutional kernel compensation algorithm [7]. In [17], an adaptive decoding algorithm was tested on a set of isometric contractions, ranging from 5% to 20% MVC, that were recorded from the tibialis anterior of eight subjects. Compared to spike trains extracted via batch decomposition, the real-time algorithm achieved an average RoA of 83%. In this work, we have achieved comparable accuracies (Figure 3) in the more challenging scenario of decoding across contraction conditions. In [24], a separate algorithm was applied to decode simulated and experimental dynamic contractions. While dynamic contractions better represent the user input of HMI applications, only pulse-to-noise ratio (PNR) was used to gauge decoder accuracy. In our study, the proposed algorithm has been directly verified using benchmark spike trains decomposed from intramuscular signals which remain the ‘gold standard’ in the field [32, 33].

### 4.1. Limitations and future work

Currently, our proposed algorithm has only been tested in a pseudo-online manner as verification of decoding accuracy against manually decomposed benchmarks necessitates offline procedures. Nonetheless, the algorithm is appropriate for real-time deployment. In this study, data windows were advanced in time increments of 100ms while the average execution time of parameter adaptation, along with spike estimation, was 57.1 *±* 14 ms. The main computational cost to the algorithm lies in the computation of the inverse covariance matrix. Despite this, prior testing has shown that omission of this step is detrimental to the overall effectiveness of the adaptive decoding algorithm. One way to reduce computational demand is to reduce the extension factor. In this work, we have used an extension factor of 16 to align with established works [8, 34]. However, past investigations suggest that lower extension factors can be employed and still retain user-intention estimation performances in HMI applications [35]. While this work focuses on verifying the accuracy of the proposed adaptive decoding algorithm against iEMG-referenced benchmarks, deployment in real-time interfacing applications is left for future works.

The adaptive decoding algorithm may not always compensate for large, sudden changes in sMUAP profiles. As show in Figure 5, when the difference between the training and test contraction conditions is significant, the adaptive algorithm may fail to converge to the correct filtering parameters. Here, the decoding algorithm is presented with an abrupt change from one isometric contraction to another, whereas, in practice, such changes occur in a continuous manner. Hence, future works will also focus on the application of adaptive decoding over experimental data pertaining to dynamic contractions.

### 5. Conclusions

In conclusion, we have developed an adaptive MU decoding algorithm that adapts to new data in realtime. Using high-confidence in-vivo-referenced benchmarks, the proposed algorithm was demonstrated to be more accurate in decoding MU activities across varying states of isometric contractions. This work therefore paves the way towards robust, real-time non-invasive neural interfacing.

## 6. Acknowledgement

This work was supported by the Academy of Finland project No. 333149, ‘Hi-Fi BiNDIng’ (I.V.), and ERC Consolidator Grant ID: 101045605 ‘INcEPTION (F.N.). We acknowledge the computational resources provided by the Aalto Science-IT project.

## References

[1] H. S. Milner-Brown, R. B. Stein, and R. Yemm. The contractile properties of human motor units during voluntary isometric contractions. The Journal of Physiology, 228(2):285, jan 1973.

[2] Jacques Duchateau and Roger M. Enoka. Human motor unit recordings: Origins and insight into the integrated motor system. Brain Research, 1409:42–61, aug 2011.

[3] C. J. de Luca, R. S. LeFever, M. P. McCue, and A. P. Xenakis. Behaviour of human motor units in different muscles during linearly varying contractions. The Journal of Physiology, 329(1):113, aug 1982.

[4] Roberto Merletti and Dario Farina. Analysis of intramuscular electromyogram signals. Philosophical Transactions of the Royal Society A: Mathematical, Physical and Engineering Sciences, 367(1887):357–368, nov 2008.

[5] J. R. Florestal, P. A. Mathieu, and K. C. McGill. Automatic decomposition of multichannel intramuscular EMG signals. Journal of Electromyography and Kinesiology, 19(1):1–9, feb 2009.

[6] Hamid R. Marateb, Silvia Muceli, Kevin C. McGill, Roberto Merletti, and Dario Farina. Robust decomposition of single-channel intramuscular EMG signals at low force levels. Journal of Neural Engineering, 8(6):066015, nov 2011.

[7] Ales Holobar and Damjan Zazula. Multichannel Blind Source Separation Using Convolution Kernel Compensation. IEEE Transactions on Signal Processing, 55(9):4487–4496, sep 2007.

[8] Francesco Negro, Silvia Muceli, Anna Margherita Castronovo, Ales Holobar, and Dario Farina. Multi-channel intramuscular and surface EMG decomposition by convolutive blind source separation. Journal of Neural Engineering, 13(2):026027, apr 2016.

[9] Maoqi Chen and Ping Zhou. A Novel Framework Based on FastICA for High Density Surface EMG Decomposition. IEEE transactions on neural systems and rehabilitation engineering : a publication of the IEEE Engineering in Medicine and Biology Society, 24(1):117–127, jan 2016.

[10] Dario Farina, Francesco Negro, Silvia Muceli, and Roger M. Enoka. Principles of motor unit physiology evolve with advances in technology. Physiology, 31(2):83–94, feb 2016.

[11] Roger M. Enoka and Dario Farina. Force steadiness: From motor units to voluntary actions. Physiology, 36(2):114–130, 2021.

[12] A. Holobar, V. Glaser, J. A. Gallego, J. L. Dideriksen, and D. Farina. Non-invasive characterization of motor unit behaviour in pathological tremor. Journal of Neural Engineering, 9(5):056011, oct 2012.

[13] Yuichi Nishikawa, Ale š Holobar, Kohei Watanabe, Tetsuya Takahashi, Hiroki Ueno, Noriaki Maeda, Hirofumi Maruyama, Shinobu Tanaka, and Allison S. Hyngstrom. Detecting motor unit abnormalities in amyotrophic lateral sclerosis using high-density surface EMG. Clinical Neurophysiology, 142:262–272, oct 2022.

[14] Dario Farina, Ivan Vujaklija, Massimo Sartori, Tamás Kapelner, Francesco Negro, Ning Jiang, Konstantin Bergmeister, Arash Andalib, Jose Principe, and Oskar C Aszmann. Man/machine interface based on the discharge timings of spinal motor neurons after targeted muscle reinnervation. Nature Biomedical Engineering, 1:25, 2017.

[15] Ales Holobar and Dario Farina. Noninvasive Neural Interfacing with Wearable Muscle Sensors: Combining Convolutive Blind Source Separation Methods and Deep Learning Techniques for Neural Decoding. IEEE Signal Processing Magazine, 38(4):103–118, jul 2021.

[16] Irene Mendez Guerra, Deren Y. Barsakcioglu, Ivan Vujaklija, Daniel Z. Wetmore, and Dario Farina. Farfield electric potentials provide access to the output from the spinal cord from wrist-mounted sensors. Journal of Neural Engineering, 19(2):026031, apr 2022.

[17] Vojko Glaser, Ales Holobar, and Damjan Zazula. Real-Time Motor Unit Identification From High-Density Surface EMG. IEEE Transactions on Neural Systems and Rehabilitation Engineering, 21(6):949–958, nov 2013.

[18] Deren Y. Barsakcioglu, Mario Bracklein, Ales Holobar, and Dario Farina. Control of Spinal Motoneurons by Feedback from a Non-invasive Real-Time Interface. IEEE Transactions on Biomedical Engineering, pages 1–1, jun 2020.

[19] Chen Chen, Yang Yu, Xinjun Sheng, Dario Farina, and Xiangyang Zhu. Simultaneous and proportional control of wrist and hand movements by decoding motor unit discharges in real time. Journal of Neural Engineering, 18(5):56010, oct 2021.

[20] Dario Farina, Marco Gazzoni, and Federico Camelia. Low-threshold motor unit membrane properties vary with contraction intensity during sustained activation with surface EMG visual feedback. Journal of Applied Physiology, 96(4):1505–1515, apr 2004.

[21] Benjamin Pasquet, Alain Carpentier, and Jacques Duchateau. Change in muscle fascicle length influences the recruitment and discharge rate of motor units during isometric contractions. Journal of Neurophysiology, 94(5):3126–3133, nov 2005.

[22] Harri Piitulainen, Timo Rantalainen, Vesa Linnamo, Paavo Komi, and Janne Avela. Innervation zone shift at different levels of isometric contraction in the biceps brachii muscle. Journal of Electromyography and Kinesiology, 19(4):667–675, aug 2009.

[23] Anderson Souza Oliveira and Francesco Negro. Neural control of matched motor units during muscle shortening and lengthening at increasing velocities. Journal of Applied Physiology, 130(6):1798–1813, jun 2021.

[24] Chen Chen, Shihan Ma, Xinjun Sheng, Dario Farina, and Xiangyang Zhu. Adaptive Real-Time Identification of Motor Unit Discharges from Non-Stationary High-Density Surface Electromyographic Signals. IEEE Transactions on Biomedical Engineering, 67(12):3501–3509, ec 2020.

[25] Aapo Hyvärinen. Fast and Robust Fixed-Point Algorithms for Independent Component Analysis. Technical Report 3, 1999.

[26] J. Thomas, Y. Deville, and S. Hosseini. Time-domain fast fixed-point algorithms for convolutive ICA. IEEE Signal Processing Letters, 13(4):228–231, apr 2006.

[27] Deren Y. Barsakcioglu and Dario Farina. A real-time surface EMG decomposition system for non-invasive human-machine interfaces. 2018 IEEE Biomedical Circuits and Systems Conference, BioCAS 2018 - Proceedings, ec 2018.

[28] A. Arturo. Leis and Vicente C. Trapani. Atlas of electromyography. Oxford University Press, 2000.

[29] Kevin C. McGill, Zoia C. Lateva, and Hamid R. Marateb. EMGLAB: An interactive EMG decomposition program. Journal of Neuroscience Methods, 149(2):121–133, ec 2005.

[30] Vojko Glaser and Ales Holobar. Motor Unit Identification From High-Density Surface Electromyograms in Repeated Dynamic Muscle Contractions. IEEE Transactions on Neural Systems and Rehabilitation Engineering, 27(1):66–75, jan 2019.

[31] Aljaz Francic and Ales Holobar. On the Reuse of Motor Unit Filters in High Density Surface Electromyograms Recorded at Different Contraction Levels. IEEE Access, 9:115227–115236, 2021.

[32] Dario Farina, Ale š Holobar, Roberto Merletti, and Roger M. Enoka. Decoding the neural drive to muscles from the surface electromyogram. Clinical Neurophysiology, 121(10):1616–1623, oct 2010.

[33] Roger M. Enoka. Physiological validation of the decomposition of surface EMG signals, jun 2019.

[34] E. Martinez-Valdes, F. Negro, C. M. Laine, D. Falla, F. Mayer, and D. Farina. Tracking motor units longitudinally across experimental sessions with highdensity surface electromyography. The Journal of Physiology, 595(5):1479–1496, mar 2017.

[35] Dennis Yeung, Francesco Negro, and I. Vujaklija. Effects of Decomposition Parameters and Estimator Type on Pseudo-online Motor Unit Based Wrist Joint Angle Prediction. Biosystems and Biorobotics, 28:371–375, 2022.

